# Impact of serum calcium levels on local and total body bone mineral density: A Mendelian randomization study and an age stratum analysis

**DOI:** 10.1101/737585

**Authors:** Jing-yi Sun, Haihua Zhang, Yan Zhang, Longcai Wang, Jin Rok Oh, Bao-liang Sun, Guiyou Liu

**Affiliations:** Department of orthopedics Wonju Severance Christian Hospital, Yonsei University Wonju College of Medicine, Wonju, Gangwon 220-701, Republic of Korea; School of Economics, Nankai University, Tianjin 300071, Tianjin, China; Department of Pathology, The Affiliated Hospital of Weifang Medical University, Weifang 261053, China; Department of Anesthesiology, The Affiliated Hospital of Weifang Medical University, Weifang 261053, China; Key Laboratory of Cerebral Microcirculation in Universities of Shandong; Department of Neurology, Second Affiliated Hospital; Shandong First Medical University & Shandong Academy of Medical Sciences, Taian 271000, Shandong, China; Department of Neurology, Xuanwu Hospital, Capital Medical University, Beijing 100053, China; Beijing Institute for Brain Disorders, Capital Medical University, Beijing, China

**Keywords:** Bone mineral density, serum calcium, Mendelian randomization

## Abstract

**Objectives:** Until recently, randomized controlled trials and meta-analyses have not demonstrated convincing conclusions regarding the association of calcium intake with bone mineral density (BMD). Until now, it remains unclear whether high serum calcium levels are causally associated with BMD. This study aimed to investigate the genetic association between serum calcium levels and BMD using a large-scale serum calcium GWAS dataset and four large-scale BMD GWAS datasets in individuals of European descent.

**Methods:** We performed a Mendelian randomization study to investigate the association of increased serum calcium levels with BMD using a large-scale serum calcium genome-wide association study (GWAS) dataset (including up to 61,079 individuals) and four large-scale BMD GWAS datasets (including minimum 4,180 individuals and maximum 142,487 individuals) regarding the total body, forearm, femoral neck, lumbar spine, and heel BMD. Here, we selected three Mendelian randomization methods including inverse-variance weighted meta-analysis (IVW), weighted median, and MR-Egger.

**Results:** In specific site analysis, we found that increased serum calcium levels could reduce BMD at forearm (OR=0.59, 95% CI: 0.36-0.95, *P*=0.029) and lumbar spine (OR=0.65, 95% CI: 0.49-0.86, *P*=0.002). We did not identify any suggestive association of genetically increased serum calcium levels with BMD of total body, femoral neck, and heel BMD. In specific age stratum analysis, we found that genetically increased serum calcium levels were statistically significantly associated with reduced total body BMD in age stratum 60 or more years (OR=0.58, 95% CI: 0.41-0.82, *P*=0.002).

**Conclusions:** We provide genetic evidence that increased serum calcium levels could not improve BMD in the general population. The elevated serum calcium levels in generally healthy populations, especially adults older than 60 years, may even reduce the BMD, and further cause osteoporosis.

## Introduction

Calcium is involved in many biological processes [1]. It is well known that calcium deficiency could cause osteoporosis [2]. Osteoporosis is a common systemic skeletal disease characterized by an increased propensity to fracture [2]. Osteoporosis could be diagnosed mainly through measurement of bone mineral density (BMD) [2]. Until now, randomized controlled trials and meta-analyses have not demonstrated convincing evidence that calcium intake (diet and supplements) could improve BMD. In fact, meta-analyses published to date have reported inconsistent conclusions regarding the association of calcium intake with BMD [3]. In 2015, Tai et al. performed a systematic review and meta-analysis of 59 randomized controlled trials [3]. They found that the increasing calcium intake from diet or supplements could produce small non-progressive increases in BMD, which are unlikely to lead to a clinically significant reduction in risk of fracture [3].

In addition to BMD, randomized controlled trials and meta-analyses published to date also have reported inconsistent conclusions regarding the association of calcium intake with osteoporosis and fracture [4-6]. In 2007, Tang et al. conducted a meta-analysis of 29 randomized trials (n=63897), which used calcium, or calcium in combination with vitamin D supplementation to prevent fracture and osteoporotic bone loss [4]. Their findings support the use of calcium, or calcium in combination with vitamin D supplementation, to improve osteoporosis in people aged 50 years or older [4]. In 2015, Bolland et al performed a systematic review of calcium intake and risk of fracture [5]. They found that dietary calcium intake was not associated with risk of fracture. Meanwhile, no clinical trial evidence shows that increasing dietary calcium intake could prevent risk of fracture [5]. There is weak and inconsistent evidence that calcium supplements prevent fractures [5]. In 2017, Zhao et al. conducted a meta-analysis of randomized clinical trials, and found that calcium supplements, vitamin D supplements, or both, were not associated with a lower risk of fractures among community-dwelling older adults compared with placebo or no treatment [6].

Due to the methodological limitations of observational studies, it is necessary to improve the causal inference through other study designs. It is reported that calcium intake (diet and supplements) especially calcium supplements could increase serum calcium levels [1, 7-9]. Until now, it remains unclear whether lifelong elevated serum calcium levels are causally associated with BMD. In recent years, large-scale genome-wide association studies (GWAS) have identified some common genetic variants and provided insight into the genetics of serum calcium levels and BMD [10-11]. These existing GWAS datasets improve the causal inference by a Mendelian randomization analysis [1, 12-17]. This method is widely used to determine the causal inferences [1, 12-17]. Here, performed a Mendelian randomization study to investigate the genetic association between serum calcium levels and BMD using a large-scale serum calcium GWAS dataset and 10 large-scale BMD GWAS datasets.

## Materials and methods

### Study Design

The Mendelian randomization is based on three principal assumptions, which have been widely described in recent studies [1, 15]. First, the genetic variants selected to be instrumental variables should be associated with the exposure (serum calcium levels) (assumption 1) [1, 15]. Second, the genetic variants should not be associated with confounders (assumption 2) [1, 15]. Third, genetic variants should affect the risk of the outcome (BMD) only through the exposure (serum calcium levels) (assumption 3) [1, 15]. The second and third assumptions are collectively known as independence from pleiotropy [15]. This study is based on the publicly available, large-scale GWAS summary datasets. All participants gave informed consent in all these corresponding original studies.

### Serum Calcium GWAS Dataset

Here, we selected genetic variants that influence serum calcium levels as the instrumental variables based on the GWAS dataset of serum calcium concentration [10]. This GWAS included 39,400 individuals from 17 population-based cohorts in discovery stage and 21,679 individuals in replication stage (N=61,079 individuals of European descent) [10]. The discovery stage and the meta-analysis of the discovery and replication stage identified 8 genetic variants to be associated with serum calcium levels with the genome-wide significance (*P* < 5.00E-08) [10]. All these 8 genetic variants were located in different genes and were not in linkage disequilibrium (Table 1) [10]. We provided more detailed information including the methods to measure serum calcium levels in **<u>eTable 1</u>**.

**Table 1.**
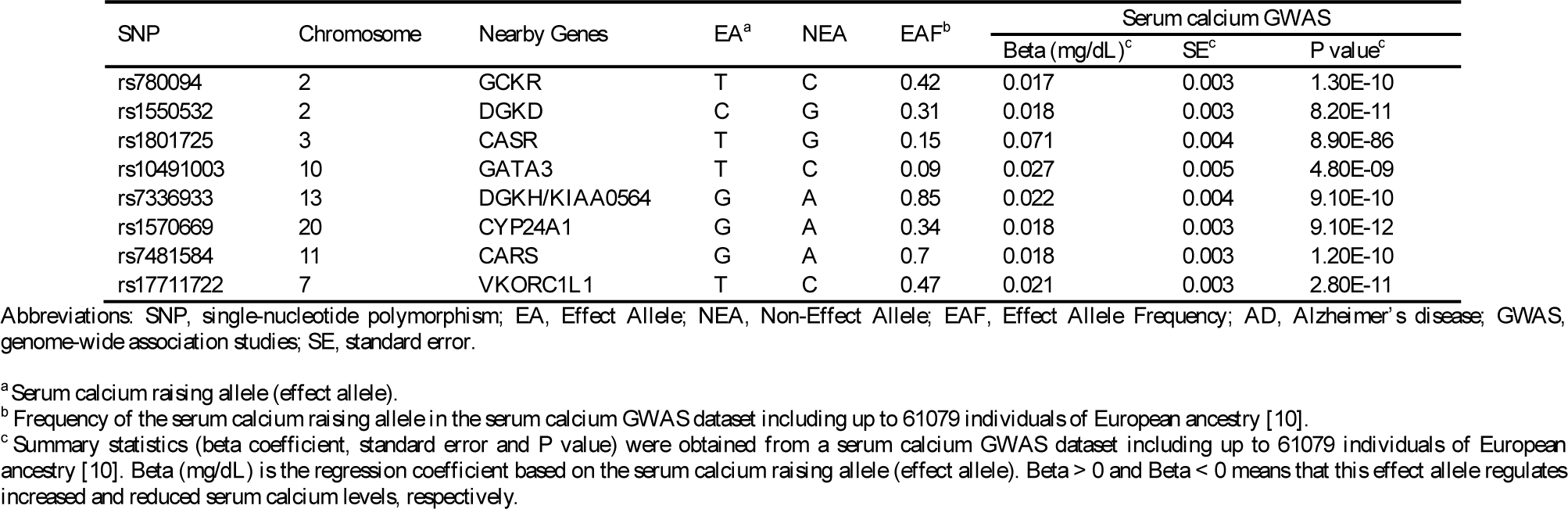
main characteristics of 8 genetic variants in serum calcium GWAS dataset

| SNP | Chromosome | Nearby Genes | EA<sup>a</sup> | NEA | EAF<sup>b</sup> | Serum calcium GWAS |  |  |
| --- | --- | --- | --- | --- | --- | --- | --- | --- |
|  |  |  |  |  |  | Beta (mg/dL)<sup>c</sup> | SE<sup>c</sup> | *P* value<sup>c</sup> |
| rs780094 | 2 | GCKR | T | C | 0.42 | 0.017 | 0.003 | 1.30E-10 |
| rs1550532 | 2 | DGKD | C | G | 0.31 | 0.018 | 0.003 | 8.20E-11 |
| rs1801725 | 3 | CASR | T | G | 0.15 | 0.071 | 0.004 | 8.90E-86 |
| rs10491003 | 10 | GATA3 | T | C | 0.09 | 0.027 | 0.005 | 4.80E-09 |
| rs7336933 | 13 | DGKH/KIAA0564 | G | A | 0.85 | 0.022 | 0.004 | 9.10E-10 |
| rs1570669 | 20 | CYP24A1 | G | A | 0.34 | 0.018 | 0.003 | 9.10E-12 |
| rs7481584 | 11 | CARS | G | A | 0.7 | 0.018 | 0.003 | 1.20E-10 |
| rs17711722 | 7 | VKORC1L1 | T | C | 0.47 | 0.021 | 0.003 | 2.80E-11 |

Abbreviations: SNP, single-nucleotide polymorphism; EA, Effect Allele; NEA, Non-Effect Allele; EAF, Effect Allele Frequency; AD, Alzheimer's disease; GWAS, genome-wide association studies; SE, standard error.
<sup>a</sup> Serum calcium raising allele (effect allele).
<sup>b</sup> Frequency of the serum calcium raising allele in the serum calcium GWAS dataset including up to 61079 individuals of European ancestry [10].
<sup>c</sup> Summary statistics (beta coefficient, standard error and *P* value) were obtained from a serum calcium GWAS dataset including up to 61079 individuals of European ancestry [10]. Beta (mg/dL) is the regression coefficient based on the serum calcium raising allele (effect allele). Beta > 0 and Beta < 0 means that this effect allele regulates increased and reduced serum calcium levels, respectively.

### BMD GWAS Datasets

Three GWAS datasets are from a large-scale meta-analysis performed by GEnetic Factors for OSteoporosis (GEFOS) Consortium and UK10K Consortia in individuals of European ancestry from the general population including BMD measured at the forearm (n=8,143), femoral neck (n=32,735) and lumbar spine (n=28,498), the sites where osteoporotic fractures are most prevalent [2]. The 4^th^ BMD dataset was measured at the heel by UK Biobank in individuals of European ancestry (n=142,487) [18]. The 5^th^ dataset is a total body-BMD GWAS including 66,628 individuals from multiple population-based cohorts across Europe (86%), America (2%), and Australia (14%) [11]. Meanwhile, single GWAS was performed in each of five age strata spanning 15 years including 0-15 years (n=11,807), 15-30 years(n=4,180), 30-45 years (n=10,062), 45-60 years(n=18,805), and 60 or more years (N=22,504) [11].

### Pleiotropy Analysis

In Mendelian randomization study, one important issue is potential violation of assumption 2 and 3 through pleiotropy occurring when a genetic instrument is associated with a study outcome through biological pathways outside the exposure of interest. Here, we performed an assessment for pleiotropy to assure that the selected genetic variants do not exert effects on BMD through biological pathways independent of serum calcium levels. We have provided more detailed information in **<u>eMethods</u>**.

### Mendelian Randomization Analysis

Here, we selected the inverse-variance weighted meta-analysis (IVW) as the main analysis method. In addition, we selected the weighted median regression and MR-Egger regression as the sensitivity analysis methods. The selection of multiple Mendelian randomization methods could examine the robustness of the estimate with each other. These methods have widely used in previous studies [1, 12-17, 19].

In order to further assess the robustness of the genetic estimates, we conducted a sensitivity analysis by sequentially removing each genetic variant from the Mendelian randomization analysis using a leave-one-out permutation analysis, which could evaluate the influence of single genetic variant on the genetic estimate. The odds ratio (OR) as well as 95% confidence interval (CI) of PD corresponds to per 0.5-mg/dL increase (about 1 standard deviation (SD)) in serum calcium levels. All analyses were conducted using the R package ‘MendelianRandomization’ [20]. The threshold for suggestive association between serum calcium levels and BMD was *P* < 0.05. The threshold of statistically significant association between serum calcium levels and BMD was a Bonferroni corrected significance *P*<0.05/10=0.005. Here, we provided more detailed information about the Mendelian randomization methods in the **<u>eMethods</u>**.

## Results

### Association of serum calcium variants with BMD

Of these 8 genetic variants associated with serum calcium levels, we successfully extracted the summary statistics for all these 8 genetic variants in each of these 10 GWAS datasets, respectively. Some of these 8 genetic variants were significantly associated with BMD at the Bonferroni corrected significance threshold (*P*<0.05/8=0.0063) (**<u>eTable 2-11</u>**).

### Pleiotropy analysis

In stage 1, rs780094 was significantly associated with known confounders at the Bonferroni corrected significance threshold (*P*<0.05/8=0.0063), as described in **eTable 12**. In brief, rs780094 variant was significantly associated with type II diabetes (*P*=1.00E-05), hip circumference (*P*=3.40E-05), waist hip ratio adjusted for BMI (*P*=1.80E-03), alcohol drinking (*P*=3.649E-09), crohns disease (*P*=2.90E-04), inflammatory bowel disease (*P*=2.24E-04), and ulcerative colitis (*P*=3.71E-03). To meet the Mendelian randomization assumptions, we excluded rs780094 variant in following analysis. In stage 2, using the remaining 7 genetic variants, MR-Egger intercept test showed no significant intercept (all *P* values > 0.05) in each of these 10 GWAS datasets (Table 2). Hence, our following analysis will be based on these 7 genetic variants.

**Table 2.**
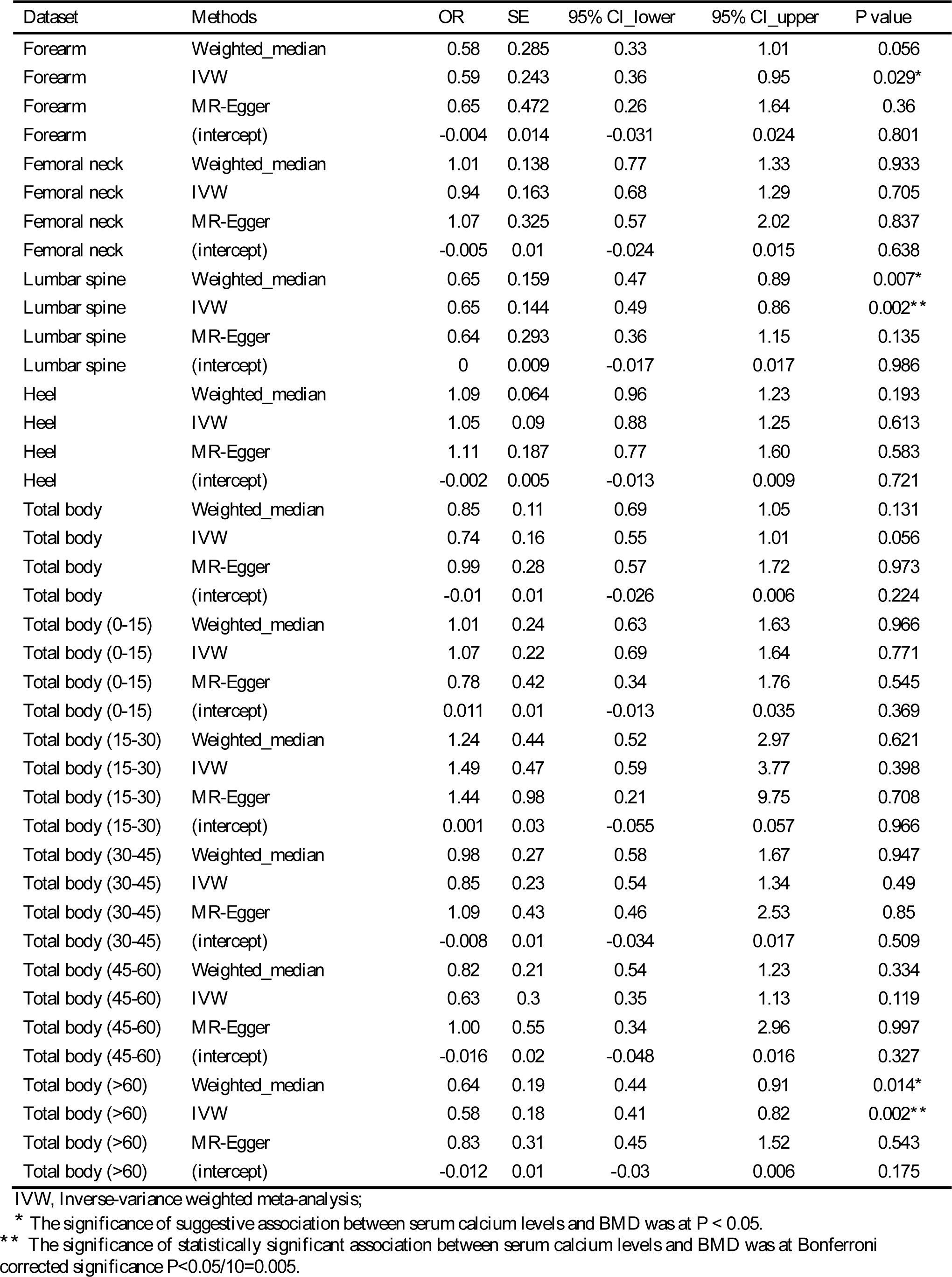
genetic association between increased serum calcium levels and BMD

### Association of serum calcium with BMD

In the forearm BMD GWAS dataset, IVW showed suggestive association between genetically increased serum calcium levels and reduced BMD (OR=0.59, 95% CI: 0.36-0.95, *P*=0.029). Interestingly, the estimates from weighted median regression, and MR-Egger were consistent with the IVW estimate in terms of direction and magnitude (Table 2). In the lumbar spine BMD GWAS dataset, IVW showed statistically significant association of genetically increased serum calcium levels with reduced BMD at a Bonferroni corrected significance *P*<0.05/10=0.005 (OR=0.65, 95% CI: 0.49-0.86, *P*=0.002). In addition, weighted median showed suggestive association of genetically increased serum calcium levels with reduced BMD (OR=0.65, 95% CI: 0.47-0.89, *P*=0.007) (Table 2). We did not identify any suggestive association of genetically increased serum calcium levels with femoral neck BMD, heel BMD, and total body-BMD, as described in Table 2.

In specific age stratum analysis, we identified no evidence of significant association of serum calcium levels with total body-BMD in four age strata including 0-15 years, 15-30 years, 30-45 years, and 45-60 years. Only in age stratum 60 or more years, IVW showed genetically increased serum calcium levels were statistically significantly associated with reduced total body-BMD at a Bonferroni corrected significance *P*<0.05/10=0.005 (OR=0.58, 95% CI: 0.41-0.82, *P*=0.002). In addition, weighted median regression showed suggestive association of genetically increased serum calcium levels with reduced total body-BMD BMD (OR=0.64, 95% CI: 0.44-0.91, *P*=1.40E-02) (Table 2). **<u>eFigure 1-10</u>** show individual genetic estimates from each of the 7 genetic variants using different methods. The leave-one-out permutation analysis further showed that the direction and precision of the genetic estimates between increased serum calcium levels and BMD remained largely unchanged using these methods.

## Discussion

It has been a long time to evaluate the association of calcium intake (diet and supplements) with osteoporosis, fracture, or BMD [3-6]. However, randomized controlled trials and meta-analyses have not demonstrated convincing evidence, but reported inconsistent conclusions regarding the association of calcium intake with osteoporosis, fracture, or BMD [3-6]. Evidence showed that calcium intake (diet and supplements) could increase serum calcium levels [1, 7-9]. Calcium supplements even could acutely increase serum calcium levels to a modest degree [1, 7-9]. Until now, it remains unclear whether elevated serum calcium levels are causally associated with BMD.

Mendelian randomization is based on the premise that the human genetic variants are randomly distributed in the population [15]. These genetic variants are largely not associated with confounders and can be used as instrumental variables to estimate the causal association of an exposure (serum calcium levels) with an outcome (BMD) [15]. Hence, Mendelian randomization could avoid some limitations of observational studies, and could be used to determine the causal inferences [1, 12-16]. Until now, the existing large-scale serum calcium and BMD GWAS datasets prompts us to investigate the potential genetic association between serum calcium and BMD by a Mendelian randomization using a large-scale serum calcium GWAS dataset and four large-scale BMD GWAS datasets.

Here, we evaluated the association of genetically increased serum calcium levels with BMD of total body and specific sites including forearm, femoral neck, lumbar spine, and heel in individuals mainly of European ancestry. The results showed that genetically increased serum calcium levels could reduce BMD at forearm and lumbar spine, but showed no association with BMD of total body, femoral neck and heel (**Table 2**). In specific age stratum analysis, our findings indicated that genetically increased serum calcium levels could only significantly reduce total body-BMD in age stratum 60 or more years in the general population. It is worth mentioning that the serum calcium levels were observed by the population-based studies including up to 61079 individuals of European descent [10]. Therefore, our conclusions reflect the effects of serum calcium levels in the general population. These conclusions may be applicable to noninstitutionalized or community-dwelling asymptomatic adults without a history of fractures. However, these conclusions may be not applicable to patients with osteoporosis, or a history of fractures, or poor serum calcium intake.

In brief, randomized controlled trials usually enrolled adults who received calcium supplementation and a concurrent comparison group that did not receive this intervention. However, randomized controlled trials did not regularly screen for the individual serum calcium status. It means that the selected individuals may have normal serum calcium levels in the beginning of the trials, or after a short time calcium supplementation. However, these individuals are still directly given a general recommendation to increase calcium supplementation. If elevated serum calcium levels are causally associated with reduced BMD in the generally healthy population, long time calcium supplementation in these individuals could not improve BMD, but even reduce the BMD, and may further cause osteoporosis. This may explain why randomized controlled trials and meta-analyses have not demonstrated convincing evidence that calcium intake (both diet and supplements) could improve BMD, and further lead to a clinically significant reduction in risk of fracture [3-6].

It is recommended that the daily calcium intake is 1000 to 1200 mg [21]. It is difficult to get this recommended amount through diet alone, so calcium supplements are widely used [21]. Until now, it remains unclear whether calcium intake from dietary sources has health advantages over supplements [22]. In the United States, about 43% of people, including about 70% of older women, take calcium supplements [23]. Hence, with the widespread use of calcium supplements, the genetic association between increased serum calcium levels and reduced BMD may have clinical and public health implications.

Our findings show that high serum calcium levels are not always better. We provide genetic evidence that high serum calcium levels could reduce BMD in the general population. If elevated serum calcium levels are causally associated with the reduced BMD in the generally healthy population, then long time calcium supplementation could not improve BMD. Therefore, our findings may explain why randomized controlled trials have not achieved convincing evidence that calcium supplements could improve BMD. Meanwhile, it is important to screen for individual serum calcium status to maximize success of randomized controlled trials. Calcium supplementation should maintain serum calcium levels at normal levels, and then may have better outcome. In addition, population-wide screening for serum calcium levels and subsequent calcium supplementation to maintain at normal levels may be a strategy for primary prevention of BMD deficiency.

This Mendelian randomization study may have several strengths. First, this study may benefit from the large-scale serum calcium GWAS dataset and BMD GWAS datasets. Second, both the serum calcium and four BMD GWAS datasets are from the European descent, which may reduce the influence on the potential association caused by the population stratification. Third, multiple independent genetic variants are taken as instruments, which may reduce the influence on the potential association caused by the linkage disequilibrium. Fourth, we performed a two-step pleiotropy analysis, and excluded one genetic variant associated with potential confounders, which meets the Mendelian randomization assumptions. Fifth, we selected multiple Mendelian randomization methods, which could reduce the risk of pleiotropy and increase the precision of the estimate. Our results are comparable with recent findings about the association of circulating serum vitamin D levels with BMD [24]. Larsson et al. found no causal association between long-term elevated circulating serum vitamin D levels and and higher BMD in generally healthy populations [24].

This Mendelian randomization study may also have several limitations. First, we provided genetic evidence that genetically increased serum calcium levels could not improve BMD, but even reduce BMD in the general population. In order to translate these genetic findings into clinical and public health implications, the potential mechanisms underlying this genetic association remain to be thoroughly evaluated. Second, we still could not completely rule out that there may be additional confounders. Until now, it is almost impossible to fully rule out pleiotropy present in any Mendelian randomization study [1, 15, 25]. Third, the GWAS dataset of serum calcium levels is from 61079 individuals of European descent [10]. We selected four BMD GWAS datasets in individuals of European ancestry to reduce the effect of population stratification [18]. In total body-BMD GWAS, most participants are from population-based cohorts of European ancestry (86%), two cohorts comprised African American individuals (2%), and four other studies included individuals with admixed background (14%) ^11^. In the original study, Medina-Gomez et al. used the LD score regression to rule out residual population stratification or cryptic relatedness ^11^. However, it could not be completely ruled out that population stratification may have had some influence on the estimate.

Fourth, the genetic association between serum calcium levels and BMD may differ by ethnicity or genetic ancestry. This genetic association should be further evaluated in other ancestries. Hence, we will further improve our work in future. Fifth, the association of serum calcium levels with additional outcomes, more clinically related, like osteoporosis and fracture could also be interesting. However, we have no access to these datasets. When these datasets are publicly available, we will further verify our findings.

In summary, we provide genetic evidence that increased serum calcium levels could not improve BMD in the general population. The lifelong elevated serum calcium levels in the generally healthy populations may even reduce the BMD.

## Acknowledgments

This work was supported by the Major State Research Development Program of China (No: 2016YFC1202302), and the National Natural Science Foundation of China (No: 61571152). We thank the GEnetic Factors for OSteoporosis (GEFOS) Consortium and UK10K Consortia for the BMD GWAS datasets. We also thank the DIAbetes Genetics Replication and Meta-analysis (DIAGRAM) Consortium, Genetic Investigation of ANthropometric Traits (GIANT) consortium, International Consortium of Blood Pressure (ICBP) consortium, Tobacco and Genetics Consortium (TGC), International Inflammatory Bowel Disease Genetics Consortium (IIBDGC), and Social Science Genetic Association Consortium (SSGAC) for other GWAS datasets.

## Author contributions

GYL and BLS conceived and initiated the project. GYL and JYS analyzed the data, drew the figures, and wrote the first draft of the manuscript. All authors contributed to the interpretation of the results and critical revision of the manuscript for important intellectual content and approved the final version of the manuscript.

## Patient consent

Obtained.

## Ethics approval

This article contains human participants collected by several studies performed by previous studies. All participants gave informed consent in all the corresponding original studies, as described in the Materials and methods. Here, our study is based on the publicly available, large-scale human GWAS summary datasets. In addition, our study does not contain any animal study.

## Reference

1. Larsson SC, Burgess S, Michaelsson K. Association of Genetic Variants Related to Serum Calcium Levels With Coronary Artery Disease and Myocardial Infarction. JAMA. 2017 Jul 25; 318(4):371–380.

2. Zheng HF, Forgetta V, Hsu YH, Estrada K, Rosello-Diez A, Leo PJ, et al. Whole-genome sequencing identifies EN1 as a determinant of bone density and fracture. Nature. 2015 Oct 1; 526(7571):112–117.

3. Tai V, Leung W, Grey A, Reid IR, Bolland MJ. Calcium intake and bone mineral density: systematic review and meta-analysis. BMJ. 2015 Sep 29; 351:h4183.

4. Tang BM, Eslick GD, Nowson C, Smith C, Bensoussan A. Use of calcium or calcium in combination with vitamin D supplementation to prevent fractures and bone loss in people aged 50 years and older: a meta-analysis. Lancet. 2007 Aug 25; 370(9588):657–666.

5. Bolland MJ, Leung W, Tai V, Bastin S, Gamble GD, Grey A, et al. Calcium intake and risk of fracture: systematic review. BMJ. 2015 Sep 29; 351:h4580.

6. Zhao JG, Zeng XT, Wang J, Liu L. Association Between Calcium or Vitamin D Supplementation and Fracture Incidence in Community-Dwelling Older Adults: A Systematic Review and Meta-analysis. JAMA. 2017 Dec 26; 318(24):2466–2482.

7. Bolland MJ, Avenell A, Baron JA, Grey A, MacLennan GS, Gamble GD, et al. Effect of calcium supplements on risk of myocardial infarction and cardiovascular events: meta-analysis. BMJ. 2010 Jul 29; 341:c3691.

8. Bolland MJ, Grey A, Avenell A, Gamble GD, Reid IR. Calcium supplements with or without vitamin D and risk of cardiovascular events: reanalysis of the Women’s Health Initiative limited access dataset and meta-analysis. BMJ. 2011 Apr 19; 342:d2040.

9. Anderson JJ, Kruszka B, Delaney JA, He K, Burke GL, Alonso A, et al. Calcium Intake From Diet and Supplements and the Risk of Coronary Artery Calcification and its Progression Among Older Adults: 10-Year Follow-up of the Multi-Ethnic Study of Atherosclerosis (MESA). J Am Heart Assoc. 2016 Oct 11; 5(10).

10. O’Seaghdha CM, Wu H, Yang Q, Kapur K, Guessous I, Zuber AM, et al. Meta-analysis of genome-wide association studies identifies six new Loci for serum calcium concentrations. PLoS Genet. 2013; 9(9):e1003796.

11. Medina-Gomez C, Kemp JP, Trajanoska K, Luan J, Chesi A, Ahluwalia TS, et al. Life-Course Genome-wide Association Study Meta-analysis of Total Body BMD and Assessment of Age-Specific Effects. Am J Hum Genet. 2018 Jan 4; 102(1):88–102.

12. Nelson CP, Hamby SE, Saleheen D, Hopewell JC, Zeng L, Assimes TL, et al. Genetically determined height and coronary artery disease. N Engl J Med. 2015 Apr 23; 372(17):1608–1618.

13. Mokry LE, Ross S, Ahmad OS, Forgetta V, Smith GD, Goltzman D, et al. Vitamin D and Risk of Multiple Sclerosis: A Mendelian Randomization Study. PLoS Med. 2015 Aug; 12(8):e1001866.

14. Manousaki D, Paternoster L, Standl M, Moffatt MF, Farrall M, Bouzigon E, et al. Vitamin D levels and susceptibility to asthma, elevated immunoglobulin E levels, and atopic dermatitis: A Mendelian randomization study. PLoS Med. 2017 May; 14(5):e1002294.

15. Emdin CA, Khera AV, Natarajan P, Klarin D, Zekavat SM, Hsiao AJ, et al. Genetic Association of Waist-to-Hip Ratio With Cardiometabolic Traits, Type 2 Diabetes, and Coronary Heart Disease. JAMA. 2017 Feb 14; 317(6):626–634.

16. Tillmann T, Vaucher J, Okbay A, Pikhart H, Peasey A, Kubinova R, et al. Education and coronary heart disease: mendelian randomisation study. BMJ. 2017 Aug 30; 358:j3542.

17. Ference BA, Kastelein JJP, Ginsberg HN, Chapman MJ, Nicholls SJ, Ray KK, et al. Association of Genetic Variants Related to CETP Inhibitors and Statins With Lipoprotein Levels and Cardiovascular Risk. JAMA. 2017 Sep 12; 318(10):947–956.

18. Kemp JP, Morris JA, Medina-Gomez C, Forgetta V, Warrington NM, Youlten SE, et al. Identification of 153 new loci associated with heel bone mineral density and functional involvement of GPC6 in osteoporosis. Nat Genet. 2017 Oct; 49(10):1468–1475.

19. Dale CE, Fatemifar G, Palmer TM, White J, Prieto-Merino D, Zabaneh D, et al. Causal Associations of Adiposity and Body Fat Distribution With Coronary Heart Disease, Stroke Subtypes, and Type 2 Diabetes Mellitus: A Mendelian Randomization Analysis. Circulation. 2017 Jun 13; 135(24):2373–2388.

20. Yavorska OO, Burgess S. MendelianRandomization: an R package for performing Mendelian randomization analyses using summarized data. Int J Epidemiol. 2017 Dec 1; 46(6):1734–1739.

21. Kern J, Kern S, Blennow K, Zetterberg H, Waern M, Guo X, et al. Calcium supplementation and risk of dementia in women with cerebrovascular disease. Neurology. 2016 Oct 18; 87(16):1674–1680.

22. Abrahamsen B. The calcium and vitamin D controversy. Ther Adv Musculoskelet Dis. 2017 May; 9(5):107–114.

23. Bailey RL, Dodd KW, Goldman JA, Gahche JJ, Dwyer JT, Moshfegh AJ, et al. Estimation of total usual calcium and vitamin D intakes in the United States. J Nutr. 2010 Apr; 140(4):817–822.

24. Larsson SC, Melhus H, Michaelsson K. Circulating Serum 25-Hydroxyvitamin D Levels and Bone Mineral Density: Mendelian Randomization Study. J Bone Miner Res. 2018 Jan 16.

25. Larsson SC, Traylor M, Malik R, Dichgans M, Burgess S, Markus HS. Modifiable pathways in Alzheimer’s disease: Mendelian randomisation analysis. BMJ. 2017 Dec 6; 359:j5375.

