## Supplementary data for "Impact of serum calcium levels on local and total body bone mineral density: A Mendelian randomization study and an age stratum analysis"

**Supplementary Online Content**

**eMethods**

**eTable 1**. Methods to measure serum calcium levels in the original study

**eTable 2**. Summary results of 8 variants with forearm BMD

**eTable 3**. Summary results of 8 variants with femoral neck BMD

**eTable 4**. Summary results of 8 variants with lumbar spine BMD

**eTable 5**. Summary results of 8 variants with heel BMD

**eTable 6**. Summary results of 8 variants with total body-BMD in overall individuals

**eTable 7**. Summary results of 8 variants with total body-BMD in individuals 0-15 years

**eTable 8**. Summary results of 8 variants with total body-BMD in individuals 15-30 years

**eTable 9**. Summary results of 8 variants with total body-BMD in individuals 30-45 years

**eTable 10**. Summary results of 8 variants with total body-BMD in individuals 45-60 years

**eTable 11**. Summary results of 8 variants with total body-BMD in individuals 60 or more years

**eTable 12**. *P* values for associations of 8 calcium-associated genetic variants with potential risk factors

**eFigure 1**. Individual genetic estimates of serum calcium levels with forearm BMD using different methods

**eFigure 2**. Individual genetic estimates of serum calcium levels with femoral neck BMD using different methods

**eFigure 3**. Individual genetic estimates of serum calcium levels with lumbar spine BMD using different methods

**eFigure 4**. Individual genetic estimates of serum calcium levels with heel BMD using different methods

**eFigure 5**. Individual genetic estimates of serum calcium levels with body-BMD in overall individuals using different methods

**eFigure 6**. Individual genetic estimates of serum calcium levels with body-BMD in individuals 0-15 years using different methods

**eFigure 7**. Individual genetic estimates of serum calcium levels with body-BMD in individuals 15-30 years using different methods

**eFigure 8**. Individual genetic estimates of serum calcium levels with body-BMD in individuals 30-45 years using different methods

**eFigure 9**. Individual genetic estimates of serum calcium levels with body-BMD in individuals 45-60 years using different methods

**eFigure 10**. Individual genetic estimates of serum calcium levels with body-BMD in individuals 60 or more years using different methods

**eMethods**

**A. Pleiotropy analysis**

In Mendelian randomization study, one important issue is potential violation of assumption 2 and 3 through pleiotropy occurring when a genetic instrument is associated with a study outcome through biological pathways outside the exposure of interest. Here, we performed an assessment for pleiotropy to assure that the selected genetic variants do not exert effects on BMD through biological pathways independent of serum calcium levels.

In stage 1, we conducted a systematic literature search to explore the potential modifiable risk factors of BMD. We identified some BMD modifiable risk factors including high blood pressure ^1^, type 2 diabetes ^2-3^, low body mass index (BMI) ^3^, smoking ^4-6^, excessive alcohol intake ^7-8^, rheumatoid arthritis ^9^, ulcerative colitis, crohns disease, or inflammatory bowel disease ^10-11^, and education ^12^. Meanwhile, lipid levels were not associated with BMD risk ^13^.

Here, we evaluated the potential pleiotropic association of each serum calcium-associated genetic variant with the potential confounders including type 2 diabetes from DIAbetes Genetics Replication and Meta-analysis (DIAGRAM) Consortium ^14^, obesity including body mass index (BMI) ^15^, waist hip ratio, waist hip ratio adjusted for BMI, waist circumference and hip circumference from Genetic Investigation of ANthropometric Traits (GIANT) consortium ^16^, systolic blood pressure (SBP) and diastolic blood pressure (DBP) from the International Consortium of Blood Pressure (ICBP) consortium ^17^, smoking behavior from the Tobacco and Genetics Consortium (TGC) (cigarettes smoked per day) ^18^, alcohol drinking (heavy vs. light) ^19^, rheumatoid arthritis ^9^, ulcerative colitis, crohns disease, or inflammatory bowel disease from International Inflammatory Bowel Disease Genetics Consortium (IIBDGC) ^10-11^, and education from Social Science Genetic Association Consortium (SSGAC) ^12, 20^. The significance threshold for the association of these 8 genetic variants with the potential confounders is a Bonferroni correction *P* < 0.05/8=0.0063.

In addition to the known confounders above, there may also be some unknown confounders. In stage 2, we selected a statistical method to evaluate the pleiotropic associations of these 8 genetic variants with other potential confounders. MR-Egger intercept test could provide an assessment of the validity of the instrumental variable assumptions, and provide a statistical test the presence of potential pleiotropy ^21-22^. More detailed information is provided in the following sections.

**B. Assumed framework of data and genetic Associations**

Suppose that we have selected J independent genetic variants () to act as instruments in a two-sample Mendelian randomization study. Here, we have successfully extracted the summary results about the associations of each genetic variant () with the risk factor (or with each risk factor for the multivariable setting) and with the outcome including the beta coefficients (,) and their standard errors (, and). We assume throughout that the parametric assumptions of linearity with no effect heterogeneity hold for the causal exposure-outcome relationship, and for the instrumental variable-exposure and instrumental variable-outcome associations for all instruments.

Initially, we consider the causal effect of a risk factor X on an outcome Y using genetic variants () that are assumed to be uncorrelated (not in linkage disequilibrium) ^23^. The association between genetic variant () and the outcome is denoted, and the association between genetic variant and the risk factor is denoted^23^. The genetic association with the outcome can be decomposed into the sum of a direct (pleiotropic) effect and an indirect (causal) effect:

where is the effect of the genetic variant on the outcome that is not mediated via the risk factor of interest, and is the causal effect of the risk factor on the outcome. A genetic variant is referred to as pleiotropic if it has associations with more than one risk factor on different causal pathways. Any such effect is included in, a genetic variant is pleiotropic if . A pleiotropic genetic variant violates the IV3 assumption, and is not a valid instrumental variable ^23^.

For a given genetic variant that meets the instrumental variable assumptions (), the causal effect of the risk factor on the outcome can be consistently estimated as a simple ratio of association estimates:

, their approximate variances

**C. Mendelian randomization analysis**

**1. Inverse-variance weighted (IVW) method**

When there are multiple genetic variants, the inverse-variance weighted (IVW) estimate is the weighted average of these causal estimates, using the inverse of their approximate variances as weights:

This estimate can be estimated using a weighted linear regression of the genetic associations with the outcome () on the genetic associations with the risk factor () using inverse variance weights () when the intercept is zero:

, weights=

Where is the residual term. If the residual standard error is set to one, this above weighted regression model, is equivalent to performing a fixed-effect meta-analysis ^23^. If the pleiotropic effects of the genetic variants are all zero ( (), in other words, if all genetic variants are valid instrumental variables), then each of the will be a consistent estimate of the causal effect, and the overall estimate (a weighted mean of the ) will be a consistent estimate of the causal effect ^23^.

**2. MR-Egger method**

The MR-Egger estimate is obtained using the same regression model the weighted linear regression described above, but allowing the intercept to be estimated as part of the analysis ^23^.

, weights=

where the parameter is the intercept, is the slope (MR-Egger estimate), and is the residual term ^23^. If the genetic variants are not pleiotropic, then the intercept term should tend to zero as the sample size increases, and the MR-Egger estimate () and the IVW estimate () are both consistent estimates of the causal effect ^23^. Alternatively, if the pleiotropic effects are independently distributed from the genetic associations with the risk factor -this is referred to as the InSIDE assumption (INstrument Strength Independent of Direct Effect) -then the MR-Egger estimate will be a consistent estimate of the causal effect as the sample size and number of genetic variants both increase ^23^.

**3. Median-based method**

The median-based method is motivated by the robustness of the median to outlying values ^24^. The median has a 50% breakdown point, meaning that median of the causal estimates obtained using each of the candidate instruments individually is a consistent estimator of the causal effect (as the sample size N tends to infinity) provided that at least half of the candidate instruments satisfy the instrumental variable assumptions ^24^. The simple median estimator is calculated as the median of the ratio estimates (). A weighted median of these causal estimates could be considered to account for differences in the precision of estimates ^24^. We assume that candidate instruments are ordered by the magnitude of their estimates (so that <<…< ), and define the weight for estimate as:

These weights are the inverse-variance weights using approximate variances. Weights are normalized so that their sum is equal to 1 ^24^. The weighted median estimate is the weighted average of the kth and (k + 1)th ratio estimates, where k is the largest integer such that the cumulative weight up to and including the kth estimate () is less than 0.5 ^24^. The estimate can be calculated by interpolation between the kth and the (k + 1)th ratio estimates:

If the intercept term is exactly equal to zero, then the MR-Egger estimate will equal the IVW estimate ^24^. The simple median estimate is the same as the weighted median estimate when all the weights are equal. For the weighted median estimate, the consistency assumption is that at least 50% of the weight in the analysis comes from valid instrumental variables ^24^.

**eTable 1**. Methods to measure serum calcium levels in the original study ^25^

| Study Name | Study Design | Genotyped sample size | Study exclusions or disease enrichment | Exclusions | Calcium Measurement + QC |
| --- | --- | --- | --- | --- | --- |
| **Discovery cohorts** |  |  |  |  |  |
| Age Gene/Environment Susceptibility Reykjavik Study (AGES) | Population based | 3664 | none | Sample exclusion criteria included sample failure, genotype mismatch with reference panel, and sex mismatch, resulting in clean genotype data on 3,219 individuals. | Serum calcium was measured using a colorimetric assay on a Hitachi 912 using a Roche Diagnostics assay . |
| ARIC | Prospective, population-based(1) | 9713 of European ancestry | none | Of the 9713 genotyped individuals of European ancestry, we excluded 658 individuals based on discrepancies with previous genotypes, disagreement between reported and genotypic sex, one randomly selected member of a pair of first-degree relatives, or outlier based on measures of average DST or more than 8 SD away on any of the first 10 principal components. | Total serum calcium was measured at ARIC visit 1 (1987-89) using a colorimetric method on a DACOS analyzer (http://www.cscc.unc.edu/aric/visit/Clinical_Chemistry_Determinations.1_10.pdf). |
| BLSA | Prospective population based study | 1230 | none | Non-European ancestry based on Eigenstrat (N=368), call rate <98.5 (N=5), sex misspecification (N=9), Missing calcium data (N=129) | Serum calcium was measured using a colorimetric assay. |
| Cohorte Lausannoise (CoLaus) | Population-based cross-sectional study | 5435 | none | Individuals with genotyping efficiency <90% were removed. Some "measured" SNPs with missing values for some individuals were imputed. As a result, the rSqHat < 1. | For each CoLaus participant a venous blood sample was collected under fasting conditions. Total serum calcium was measured by O-cresolphtalein (2.1% –1.5% maximum inter and intra-batch CVs); albumin was measured by bromocresol green (2.5% – 0.4%). |
| CROATIA-Vis | Family-based, cross-sectional study in an isolated population | 924 | none | Exclusions: sample call rate<95% | Serum calcium was measured using a colorimetric assay. |
| CROATIA-Korcula | Family-based, cross-sectional study in an isolated population | 899 | none | Exclusions: sample call rate<97% | Serum calcium was measured using a colorimetric assay. |
| CROATIA-Split | Population-based, cross-sectional study | 499 | none | Exclusions: sample call rate<97% | Serum calcium was measured using a colorimetric assay. |
| Framingham Heart Study (FHS) | Prospective family-based | 9300 | none | Of the 9,274 participants who underwent genotyping, we made the following exclusions: sample call rate <97% (n=666), genotype heterozygosity > 5 standard deviations, and ambiguous family data (n=127). This resulted in a total of 8,481 genotyped individuals. | Serum calcium was measured using a colorimetric assay. |
| HABC | prospective cohort study | 1663 | none | none | Serum calcium was measured using a colorimetric assay. |
| InCHIANTI | Prospective population based study | 1231 | none | Ambiguous family data (N=4), call rate < 98.5% (N=12), sex misspecification (N=1), heterozygosity>0.3 (N=4), Missing calcium data (N=6) | Serum calcium was measured using a colorimetric assay. |
| Lothian Birth Cohort 1936 (LBC1936) | Population based birth cohort | 1005 | none | Individuals with a disagreement between genetic and reported gender were removed (n = 12). Relatedness between subjects was investigated and, for any related pair of individuals, one was removed [PI_HAT (proportion of IBD) > 0.25, n = 8). Samples with a call rate ≤ 0.95 (n = 16), and those showing evidence of non-European descent by multidimensional scaling, were also removed (n = 1). | Serum calcium was measured using a colorimetric assay. |
| London Life Sciences Population (LOLIPOP) study | | |  |  |  |
| LOLIPOP EW A | Population based prospective cohort study | 878 | none | excluded samples of duplicates, contaminated samples, call rate <%95, and the same samples apeared in EW610 | Serum calcium was measured using standard approach |
| LOLIPOP EW P | Population based prospective cohort study | 1005 | none | excluded samples of duplicates, contaminated samples, call rate <%95, and the same samples apeared in EW610 | Serum calcium was measured using standard approach |
| LOLIPOP EW610 | Population based prospective cohort study | 945 | none | excluded samples of duplicates, call rate <%95, relatedness, gender discrepancy, ethnic outliers, and imcomplete clinical data | Serum calcium was measured using standard approach |
| Ogliastra Genetic Park - Talana Study | Population-based study with pedigree information | 860 | none |  | Calcium levels were determined with an automated Targa BT-3000 Chemistry Analyser ( ml/dL) |
| ORCADES | Family-based, cross-sectional study in an isolated population | 889 | none | Exclusions: sample call rate<97% | Serum calcium was measured using a colorimetric assay. |
| SHIP | Prospective population-based study | 4081 | none | Excluded arrays with CallRate < 92%, duplicate samples (by estimated IBD) and individuals with reported/genotyped gender mismatch | Non-fasting blood samples were drawn from the cubital vein in the supine position between 7.00 a.m. and 7.00 p.m. The measurement of total calcium concentration was performed immediately after blood withdrawal. Samples were analysed on the Hitachi 911 by o-cresolphthalein complexone colorimetry (Boehringer Mannheim, Germany). The internal quality controls were analyzed daily. During the course of the study the inter as well as the intra-assay coefficient of variation was <5%. In addition, the laboratory takes part quarterly in the official national German external proficiency testing programs and fulfilled the requirements. |
| The Rotterdam Study (RS) | Prospective population based study | 5974 | NA | Any samples with a call rate below 97.5%, excess autosomal heterozygosity >0.336 (~FDR <0.1%), mismatch between called and phenotypic gender, or if there were outliers identified by the IBS clustering analysis (see below) with >3 standard deviations from population mean or IBS probabilities >97% were excluded from the analysis | Serum calcium was measured at baseline visit in the Rotterdam Study with a colorimetric detection assay using the Hitachi 917 (Roche, Mannheim, Germany). |
| The Cardiovascular Health Study (CHS) | Prospective, population-based | 3,329 CHS Caucasian participants | A total of 1908 persons were excluded from the GWAS study sample due to the presence at study baseline of coronary heart disease, congestive heart failure, peripheral vascular disease, valvular heart disease, stroke or transient ischemic attack or lack of available DNA. | The present report is based upon genotyping results from 3,329 CHS Caucasian participants, who were free of clinical cardiovascular disease at baseline, consented to genetic testing, and had DNA available for genotyping. Genotypes were called using the Illumina BeadStudio software. Genotyping was successful in 3,291 persons. | Serum calcium has been measured in unit mg/dL |
| **Replication cohorts** |  |  |  |  |  |
| Bus Santé study | Cross-sectional population-based study | 5622 | none | Of the 5,622 participants who underwent genotyping, genotyping was unsuccessfull for 2.7% (N=151). This resulted in a total of 5,471 genotyped individuals. Analyses were restriced to Caucasians. Caucasian was defined as self-reported citizenship corresponding to South/North America, Europe, and Australia regions | Serum calcium was measured using a colorimetric assay using Arsenazo-III reactive (Architect CI4100®, Abbott). Coefficient of variation (CV) = 5.4% |
| INGI-Carlantino-Project | Isolated population | 679 | none | we made the following exclusions: sample call rate <97% | Serum calcium was measured using a colorimetric assay. |
| INGI-FVG-Project | Isolated population | 1471 | none | we made the following exclusions: sample call rate <97% | Serum calcium was measured using a colorimetric assay. |
| INGI-CILENTO | Cross-sectional population based | 1147 | none | none | Serum calcium was measured using a colorimetric assay. |
| KORA F3 Study (Cooperative Health Research in the Region of Augsburg) | Population-based | 1643 | none | 3, because of no available information on serum calcium. This resulted in a total of 1640 individuals | Serum calcium was measured using a colorimetric assay. |
| KORA F4 Study | Population-based | 1814 | none | 5, because of no available  information on serum calcium. This resulted in a total of 1809 individuals | Serum calcium was measured using a colorimetric assay. |
| LURIC Study | Case-control | 3032 |  | sample call rate <95%, gender ambiguity, relatedness. This resulted in a total of 2927 genotyped individuals. | o-Kresolphthalein-complexon, CA/Hitachi 717, Roche, Germany |
| PIVUS | Prospective cohort | 958 | none | Sample call rate <95%, genotype heterozygosity > +-3 standard deviations, gender discordance, and duplicates. | Reference method at Uppsala University Hospital |
| SHIP-Trend | Prospective population-based study | 986 | none | Excluded arrays with CallRate < 94%, duplicate samples (by estimated IBD) and individuals with reported/genotyped gender mismatch | Blood samples in SHIP-Trend were taken, while subjects were on random salt diet and under regular medication, from fasting subjects in supine position. Serum calcium has been measured by photometric rocedure of bichromatic endpoint measurement on an Dimension Vista system (Dade Behring, Eschborn, Germany). The analytical measurement range was: 1.25-3.75 mmol/L. During the course of the study the inter-assay coefficient of variation was < 5%. |
| TwinsUK | Twin Study | 3965 | none | Of the 5,654 participants who underwent genotyping, we made the following exclusions: >3 standard deviations and missing informations. This resulted in a total of 3965 genotyped individuals. | Assays for serum calcium was performed using standard laboratory procedures. The test was performed on a 950 Vitros analyser (Ortho-Clinical Diagnostics; Johnson and Johnson, Rochester, NY, U.S.A.). |
| The BRItish Genetics of HyperTension (BRIGHT) study | Hypertensive cases from the BRIGHT study resource. | 2000 | BMI>35  diabetes, secondary hypertension or a co-existing illness. | Of 2000 cases typed, we excluded individuals if they had >3% missing data or evidence of non-European ancestry under eigenstrat analysis, n=277. | Serum calcium measures were performed on non-fasting samples by the Clinical Biochemistry Unit at the University of Glasgow |

**eTable 2**. Summary results of 8 variants with forearm BMD ^26^

| SNP | EA | NEA | EAF | Beta | SE | *P* value |
| --- | --- | --- | --- | --- | --- | --- |
| rs780094 | T | C | 0.352745 | -0.03936 | 0.015928 | 0.015404 |
| rs1550532 | C | G | 0.273031 | -0.00349 | 0.016812 | 0.838881 |
| rs17711722 | T | C | 0.44471 | -0.01029 | 0.019344 | 0.601861 |
| rs10491003 | T | C | 0.099364 | 0.037436 | 0.027261 | 0.178123 |
| rs7481584 | A | G | 0.316229 | 0.018214 | 0.01709 | 0.29602 |
| rs7336933 | A | G | 0.140255 | 0.036037 | 0.021514 | 0.10051 |
| rs1570669 | G | A | 0.393317 | -0.01718 | 0.016063 | 0.294286 |
| rs17251221 | G | A | 0.116309 | -0.03846 | 0.022234 | 0.089905 |

SNP, single-nucleotide polymorphism; EA, Effect Allele; NEA, Non-Effect Allele; EAF, Effect Allele Frequency; SE, standard error. Beta > 0 and Beta < 0 means that this effect allele regulates increased and reduced BMD, respectively.

**eTable 3**. Summary results of 8 variants with femoral neck BMD ^26^

| SNP | EA | NEA | EAF | Beta | SE | *P* value |
| --- | --- | --- | --- | --- | --- | --- |
| rs780094 | T | C | 0.352745 | -0.01504 | 0.007663 | 0.054802 |
| rs1550532 | C | G | 0.273031 | -0.01553 | 0.008436 | 0.071614 |
| rs17711722 | T | C | 0.44471 | 0.001282 | 0.009048 | 0.889706 |
| rs10491003 | T | C | 0.099364 | 0.001175 | 0.01323 | 0.930717 |
| rs7481584 | A | G | 0.316229 | 0.007112 | 0.008466 | 0.411069 |
| rs7336933 | A | G | 0.140255 | 0.021365 | 0.010577 | 0.048125 |
| rs1570669 | G | A | 0.393317 | 0.015003 | 0.007801 | 0.059892 |
| rs17251221 | G | A | 0.116309 | 0.000755 | 0.010654 | 0.944704 |

SNP, single-nucleotide polymorphism; EA, Effect Allele; NEA, Non-Effect Allele; EAF, Effect Allele Frequency; SE, standard error. Beta > 0 and Beta < 0 means that this effect allele regulates increased and reduced BMD, respectively.

**eTable 4**. Summary results of 8 variants with lumbar spine BMD ^26^

| SNP | EA | NEA | EAF | Beta | SE | *P* value |
| --- | --- | --- | --- | --- | --- | --- |
| rs780094 | T | C | 0.352745 | -0.02208 | 0.008935 | 0.015812 |
| rs1550532 | C | G | 0.273031 | -0.02619 | 0.009463 | 0.006881 |
| rs17711722 | T | C | 0.44471 | -0.00223 | 0.009943 | 0.826826 |
| rs10491003 | T | C | 0.099364 | 0.009478 | 0.015322 | 0.545781 |
| rs7481584 | A | G | 0.316229 | 0.005259 | 0.0098 | 0.600209 |
| rs7336933 | A | G | 0.140255 | 0.013639 | 0.012272 | 0.277718 |
| rs1570669 | G | A | 0.393317 | -0.00236 | 0.009082 | 0.799414 |
| rs17251221 | G | A | 0.116309 | -0.03363 | 0.012377 | 0.007978 |

SNP, single-nucleotide polymorphism; EA, Effect Allele; NEA, Non-Effect Allele; EAF, Effect Allele Frequency; SE, standard error. Beta > 0 and Beta < 0 means that this effect allele regulates increased and reduced BMD, respectively.

**eTable 5**. Summary results of 8 variants with heel BMD ^27^

| SNP | EA | NEA | EAF | Beta | SE | *P* value |
| --- | --- | --- | --- | --- | --- | --- |
| rs780094 | T | C | 0.384449 | -0.00854 | 0.003423 | 5.80E-03 |
| rs1550532 | C | G | 0.316341 | 0.002104 | 0.003591 | 7.40E-01 |
| rs1801725 | G | T | 0.86876 | -0.00568 | 0.00497 | 2.90E-01 |
| rs17711722 | C | T | 0.549161 | -0.00865 | 0.003349 | 1.50E-02 |
| rs10491003 | C | T | 0.907363 | 0.004981 | 0.00579 | 4.40E-01 |
| rs7481584 | G | A | 0.715199 | -0.0093 | 0.003714 | 1.80E-02 |
| rs7336933 | G | A | 0.847918 | -0.00725 | 0.004701 | 2.30E-01 |
| rs1570669 | A | G | 0.655904 | -0.00312 | 0.003563 | 6.30E-01 |

SNP, single-nucleotide polymorphism; EA, Effect Allele; NEA, Non-Effect Allele; EAF, Effect Allele Frequency; SE, standard error. Beta > 0 and Beta < 0 means that this effect allele regulates increased and reduced BMD, respectively.

**eTable 6**. Summary results of 8 variants with total body-BMD in all individuals ^28^

| SNP | EA | NEA | EAF | Beta | SE | *P* value |
| --- | --- | --- | --- | --- | --- | --- |
| rs1550532 | t | c | 0.3772 | -0.0301 | 0.0058 | 2.54E-07 |
| rs7336933 | c | g | 0.3131 | -0.025 | 0.0061 | 4.09E-05 |
| rs1801725 | a | g | 0.1505 | 0.0239 | 0.0079 | 0.00265 |
| rs10491003 | t | g | 0.1282 | -0.0103 | 0.0083 | 0.2157 |
| rs17711722 | t | c | 0.0941 | -0.008 | 0.0098 | 0.418 |
| rs7481584 | t | c | 0.417 | -0.0051 | 0.0072 | 0.4804 |
| rs1570669 | a | g | 0.2985 | 0.0014 | 0.0063 | 0.83 |
| rs780094 | a | g | 0.6486 | -9.00E-04 | 0.006 | 0.8819 |

SNP, single-nucleotide polymorphism; EA, Effect Allele; NEA, Non-Effect Allele; EAF, Effect Allele Frequency; SE, standard error. Beta > 0 and Beta < 0 means that this effect allele regulates increased and reduced BMD, respectively.

**eTable 7**. Summary results of 8 variants with total body-BMD in individuals 0-15 years ^28^

| SNP | EA | NEA | EAF | Beta | SE | *P* value |
| --- | --- | --- | --- | --- | --- | --- |
| rs1550532 | c | g | 0.299 | -0.0226 | 0.0141 | 0.1089 |
| rs780094 | t | c | 0.3739 | -0.0208 | 0.0136 | 0.1257 |
| rs1570669 | a | g | 0.6462 | -0.0175 | 0.0135 | 0.1938 |
| rs17711722 | t | c | 0.4124 | 0.0159 | 0.0173 | 0.3569 |
| rs7336933 | a | g | 0.1487 | -0.0156 | 0.0181 | 0.3874 |
| rs7481584 | a | g | 0.2832 | -0.0113 | 0.0144 | 0.4302 |
| rs1801725 | t | g | 0.133 | -0.0086 | 0.0192 | 0.6531 |
| rs10491003 | t | c | 0.0942 | 0.0055 | 0.0222 | 0.8056 |

SNP, single-nucleotide polymorphism; EA, Effect Allele; NEA, Non-Effect Allele; EAF, Effect Allele Frequency; SE, standard error. Beta > 0 and Beta < 0 means that this effect allele regulates increased and reduced BMD, respectively.

**eTable 8**. Summary results of 8 variants with total body-BMD in individuals 15-30 years ^28^

| SNP | EA | NEA | EAF | Beta | SE | *P* value |
| --- | --- | --- | --- | --- | --- | --- |
| rs10491003 | t | c | 0.0936 | 0.0737 | 0.0396 | 0.06261 |
| rs780094 | t | c | 0.3826 | -0.043 | 0.0231 | 0.06292 |
| rs7481584 | a | g | 0.304 | 0.0419 | 0.0249 | 0.09284 |
| rs7336933 | a | g | 0.1459 | -0.0506 | 0.0323 | 0.1171 |
| rs1570669 | a | g | 0.6578 | 0.0257 | 0.0241 | 0.2855 |
| rs1550532 | c | g | 0.3173 | -0.0238 | 0.0239 | 0.3201 |
| rs1801725 | t | g | 0.1258 | 0.0165 | 0.0342 | 0.6292 |
| rs17711722 | t | c | 0.4054 | -0.0021 | 0.0293 | 0.9416 |

SNP, single-nucleotide polymorphism; EA, Effect Allele; NEA, Non-Effect Allele; EAF, Effect Allele Frequency; SE, standard error. Beta > 0 and Beta < 0 means that this effect allele regulates increased and reduced BMD, respectively.

**eTable 9**. Summary results of 8 variants with total body-BMD in individuals 30-45 years ^28^

| SNP | EA | NEA | EAF | Beta | SE | *P* value |
| --- | --- | --- | --- | --- | --- | --- |
| rs780094 | t | c | 0.3822 | -0.0321 | 0.0149 | 0.03112 |
| rs7336933 | a | g | 0.1418 | 0.0403 | 0.0211 | 0.05573 |
| rs1550532 | c | g | 0.3162 | -0.0152 | 0.0156 | 0.329 |
| rs10491003 | t | c | 0.0978 | -0.0136 | 0.0255 | 0.5923 |
| rs7481584 | a | g | 0.3188 | -0.0076 | 0.016 | 0.6352 |
| rs1570669 | a | g | 0.6417 | -0.0035 | 0.0154 | 0.8183 |
| rs17711722 | t | c | 0.413 | -0.0027 | 0.0178 | 0.8804 |
| rs1801725 | t | g | 0.1244 | 5.00E-04 | 0.0213 | 0.9825 |

SNP, single-nucleotide polymorphism; EA, Effect Allele; NEA, Non-Effect Allele; EAF, Effect Allele Frequency; SE, standard error. Beta > 0 and Beta < 0 means that this effect allele regulates increased and reduced BMD, respectively.

**eTable 10**. Summary results of 8 variants with total body-BMD in individuals 45-60 years ^28^

| SNP | EA | NEA | EAF | Beta | SE | *P* value |
| --- | --- | --- | --- | --- | --- | --- |
| rs780094 | t | c | 0.3793 | -0.0459 | 0.0108 | 2.11E-05 |
| rs1550532 | c | g | 0.3177 | -0.0376 | 0.0112 | 0.000815 |
| rs7336933 | a | g | 0.1467 | 0.0439 | 0.015 | 0.003496 |
| rs10491003 | t | c | 0.0919 | -0.0379 | 0.019 | 0.04557 |
| rs7481584 | a | g | 0.3028 | -0.0108 | 0.0118 | 0.3621 |
| rs1801725 | t | g | 0.122 | -0.0121 | 0.0156 | 0.4355 |
| rs17711722 | t | c | 0.4221 | -0.0063 | 0.0136 | 0.6461 |
| rs1570669 | a | g | 0.644 | 0.0045 | 0.0111 | 0.6886 |

SNP, single-nucleotide polymorphism; EA, Effect Allele; NEA, Non-Effect Allele; EAF, Effect Allele Frequency; SE, standard error. Beta > 0 and Beta < 0 means that this effect allele regulates increased and reduced BMD, respectively.

**eTable 11**. Summary results of 8 variants with total body-BMD in individuals 60 or more years ^28^

| SNP | EA | NEA | EAF | Beta | SE | *P* value |
| --- | --- | --- | --- | --- | --- | --- |
| rs1550532 | c | g | 0.3165 | -0.0299 | 0.0106 | 0.004801 |
| rs7336933 | a | g | 0.1582 | 0.0299 | 0.0134 | 0.02512 |
| rs1801725 | t | g | 0.1312 | -0.0268 | 0.0142 | 0.05907 |
| rs780094 | t | c | 0.3725 | -0.0189 | 0.01 | 0.06 |
| rs7481584 | a | g | 0.2957 | 0.0151 | 0.0108 | 0.1624 |
| rs17711722 | t | c | 0.416 | -0.0159 | 0.0123 | 0.1952 |
| rs1570669 | a | g | 0.6548 | 0.0014 | 0.0104 | 0.8942 |
| rs10491003 | t | c | 0.0944 | -5.00E-04 | 0.0167 | 0.9743 |

SNP, single-nucleotide polymorphism; EA, Effect Allele; NEA, Non-Effect Allele; EAF, Effect Allele Frequency; SE, standard error. Beta > 0 and Beta < 0 means that this effect allele regulates increased and reduced BMD, respectively.

**eTable 12**. *P* Values for Associations of 8 Calcium-Associated Genetic Variants with potential risk factors

| SNP | DBP | SBP | Type 2 diabetes | Smoking | Alcohol drinking | BMI | Hip circumference | Waist circumference | Waist hip ratio | Waist hip ratio adjusted for BMI | Crohns disease | Inflammatory bowel disease | Ulcerative colitis | Rheumatoid arthritis |
| --- | --- | --- | --- | --- | --- | --- | --- | --- | --- | --- | --- | --- | --- | --- |
| rs10491003 | 0.921 | 0.956 | 0.83 | 0.9987 | 0.6499 | 0.5716 | 0.69 | 0.17 | 0.3021 | 0.044 | 0.7826 | 0.9726 | 0.8908 | 0.23 |
| rs1550532 | 0.118 | 0.113 | 0.58 | 0.4318 | 0.386 | 0.5101 | 0.95 | 0.59 | 0.6247 | 0.91 | 0.09032 | 0.01167 | 0.06552 | 0.81 |
| rs1570669 | 0.982 | 0.962 | 0.097 | 0.487 | 0.4071 | 0.337 | 0.5 | 0.41 | 0.3787 | 0.67 | 0.6013 | 0.6576 | 0.9271 | 0.088 |
| rs17711722 | 0.563 | 0.986 | 0.46 | 0.2361 | 0.6723 | 0.2342 | 0.091 | 0.13 | 0.6016 | 0.63 | 0.2167 | 0.2346 | 0.6449 | NA |
| rs1801725 | 0.969 | 0.261 | 0.35 | 0.09868 | 0.7526 | 0.4853 | 0.29 | 0.51 | 0.4825 | 0.26 | 0.03717 | 0.8119 | 0.01892 | 0.059 |
| rs7336933 | 0.0553 | 0.269 | 0.87 | 0.457 | 0.8629 | 0.2971 | 0.81 | 0.91 | 0.8309 | 0.78 | 0.5519 | 0.5409 | 0.8712 | 0.77 |
| rs7481584 | 0.397 | 0.684 | 0.015 | 0.5377 | 0.393 | 0.3346 | 0.11 | 0.064 | 0.2543 | 0.24 | 0.6283 | 0.5626 | 0.9089 | 0.83 |
| rs780094 | 0.544 | 0.313 | **1.00E-05** | 0.2184 | **3.649E-09** | 0.05802 | **3.40E-05** | 0.015 | 0.02108 | **1.80E-03** | **2.90E-04** | **2.24E-04** | **3.71E-03** | 0.45 |

In stage 1, we conducted a systematic literature search to explore the potential modifiable risk factors of BMD. We identified some BMD modifiable risk factors including high blood pressure ^1^, type 2 diabetes ^2-3^, low body mass index (BMI) ^3^, smoking ^4-6^, excessive alcohol intake ^7-8^, rheumatoid arthritis ^9^, ulcerative colitis, crohns disease, or inflammatory bowel disease ^10-11^, and education ^12^. Meanwhile, lipid levels were not associated with BMD risk ^13^.

Here, we evaluated the potential pleiotropic association of each serum calcium-associated genetic variant with the potential confounders including type 2 diabetes from DIAbetes Genetics Replication and Meta-analysis (DIAGRAM) Consortium ^14^, obesity including body mass index (BMI) ^15^, waist hip ratio, waist hip ratio adjusted for BMI, waist circumference and hip circumference from Genetic Investigation of ANthropometric Traits (GIANT) consortium ^16^, systolic blood pressure (SBP) and diastolic blood pressure (DBP) from the International Consortium of Blood Pressure (ICBP) consortium ^17^, smoking behavior from the Tobacco and Genetics Consortium (TGC) (cigarettes smoked per day) ^18^, alcohol drinking (heavy vs. light) ^19^, rheumatoid arthritis ^9^, ulcerative colitis, crohns disease, or inflammatory bowel disease from International Inflammatory Bowel Disease Genetics Consortium (IIBDGC) ^10-11^, and education from Social Science Genetic Association Consortium (SSGAC) ^12, 20^. The significance threshold for the association of these 8 genetic variants with the potential confounders is a Bonferroni correction *P* < 0.05/8=0.0063.

**eFigure 1**. Individual genetic estimates of serum calcium levels with forearm BMD using different methods

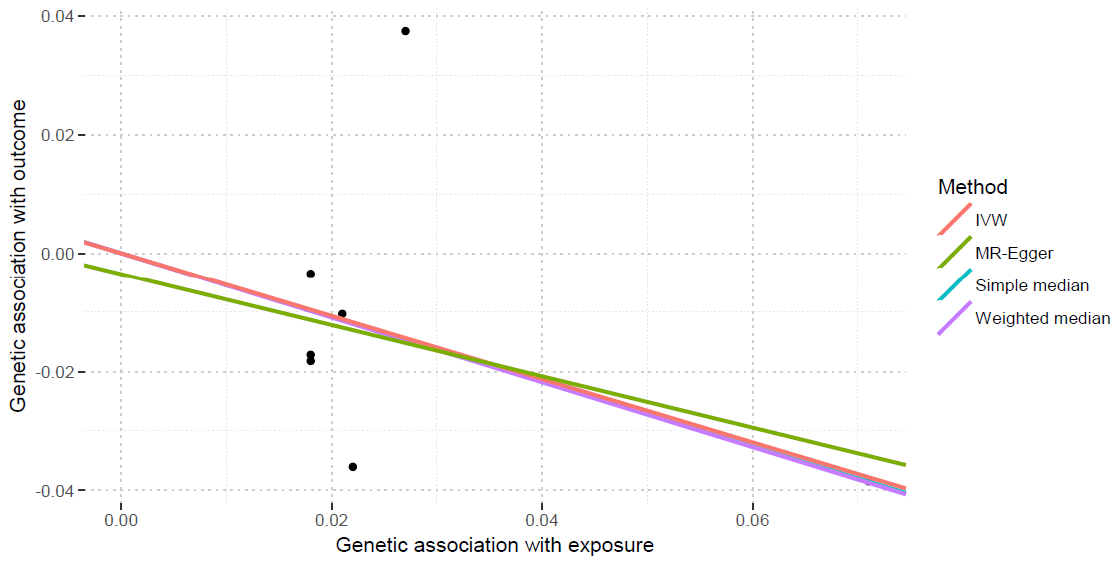

This scatter plot show individual causal estimates from each of 7 genetic variants associated with serum calcium levels on the x-axis and BMD on the y-axis. The continuous line represents the causal estimate of serum calcium levels on BMD.

**eFigure 2**. Individual genetic estimates of serum calcium levels with Femoral neck BMD using different methods

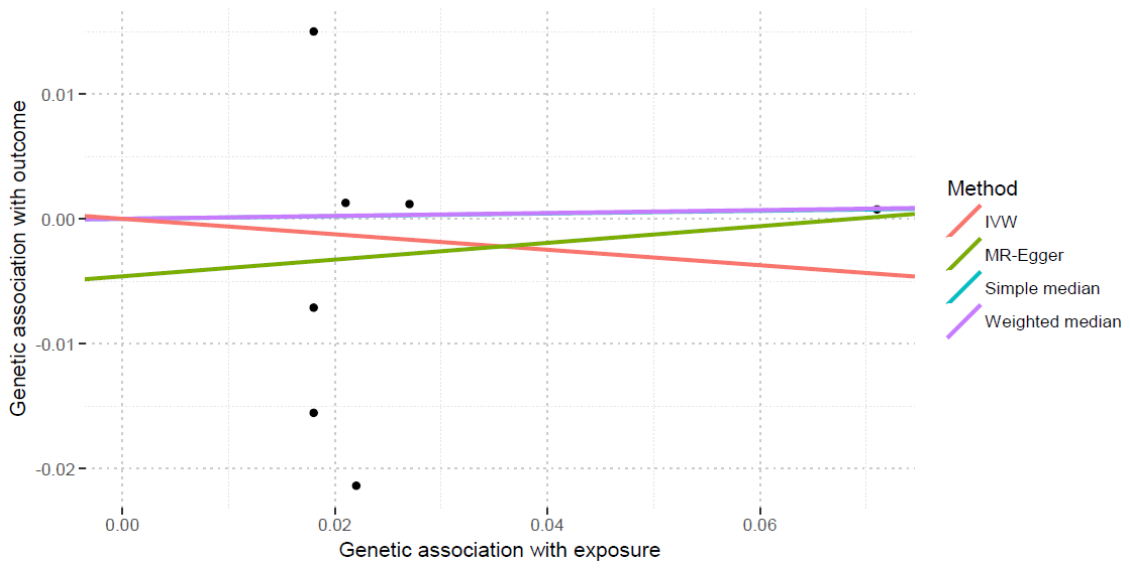

This scatter plot show individual causal estimates from each of 7 genetic variants associated with serum calcium levels on the x-axis and BMD on the y-axis. The continuous line represents the causal estimate of serum calcium levels on BMD.

**eFigure 3**. Individual genetic estimates of serum calcium levels with Lumbar spine BMD using different methods

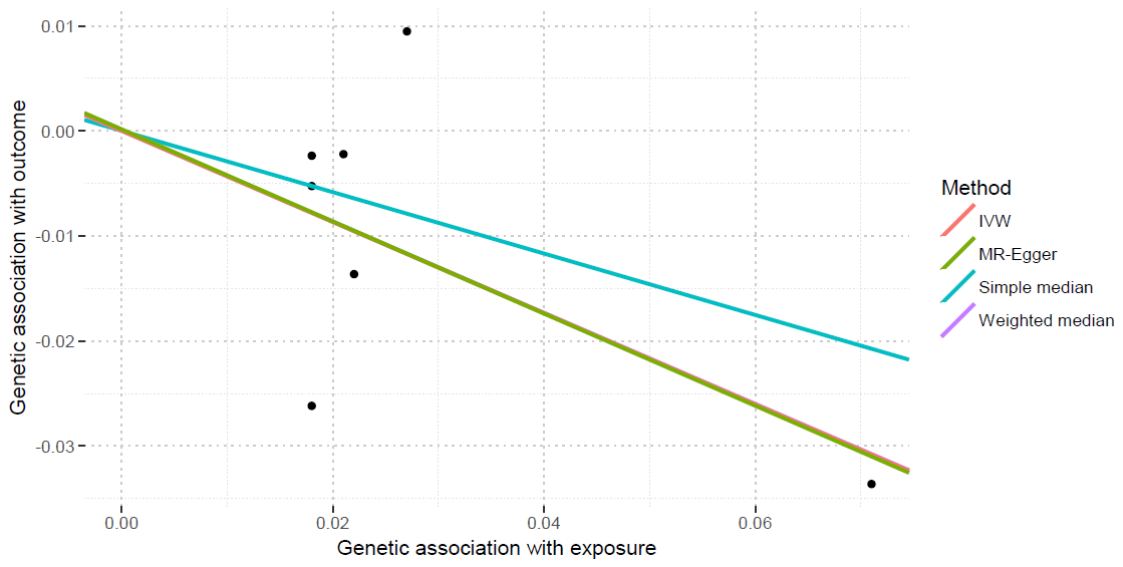

This scatter plot show individual causal estimates from each of 7 genetic variants associated with serum calcium levels on the x-axis and BMD on the y-axis. The continuous line represents the causal estimate of serum calcium levels on BMD.

**eFigure 4**. Individual genetic estimates of serum calcium levels with hell BMD using different methods

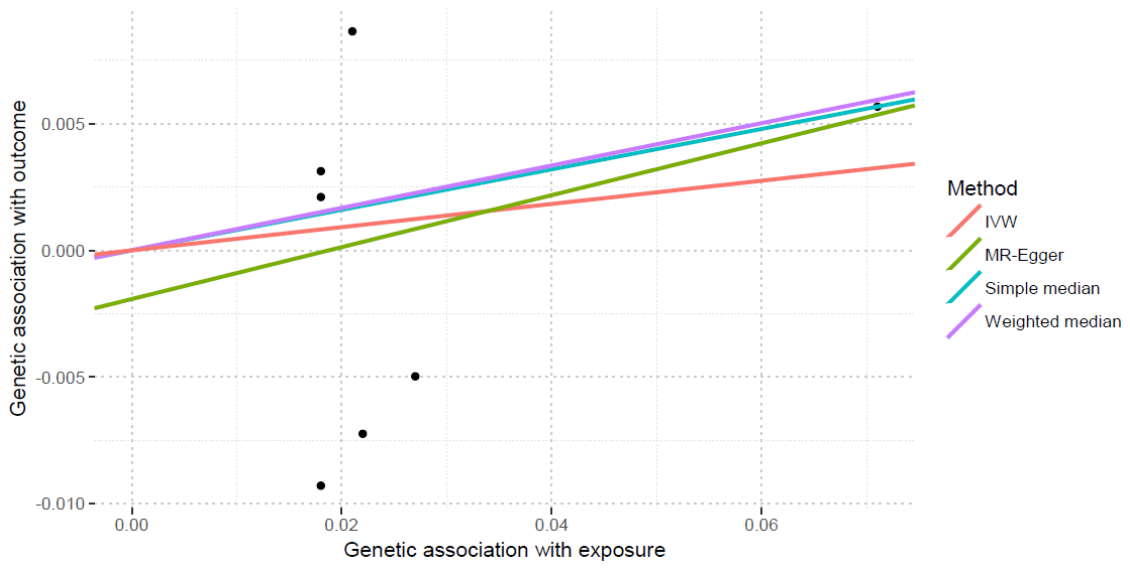

This scatter plot show individual causal estimates from each of 7 genetic variants associated with serum calcium levels on the x-axis and BMD on the y-axis. The continuous line represents the causal estimate of serum calcium levels on BMD.

**eFigure 5**. Individual genetic estimates of serum calcium levels with body-BMD in all individuals using different methods

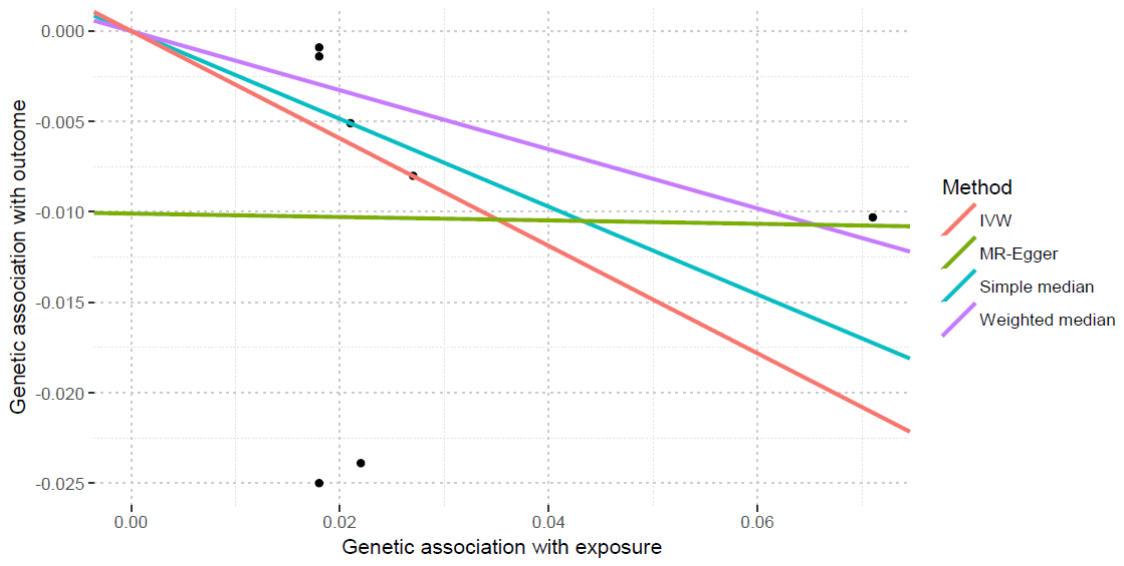

This scatter plot show individual causal estimates from each of 7 genetic variants associated with serum calcium levels on the x-axis and BMD on the y-axis. The continuous line represents the causal estimate of serum calcium levels on BMD.

**eFigure 6**. Individual genetic estimates of serum calcium levels with body-BMD in individuals 0-15 years using different methods

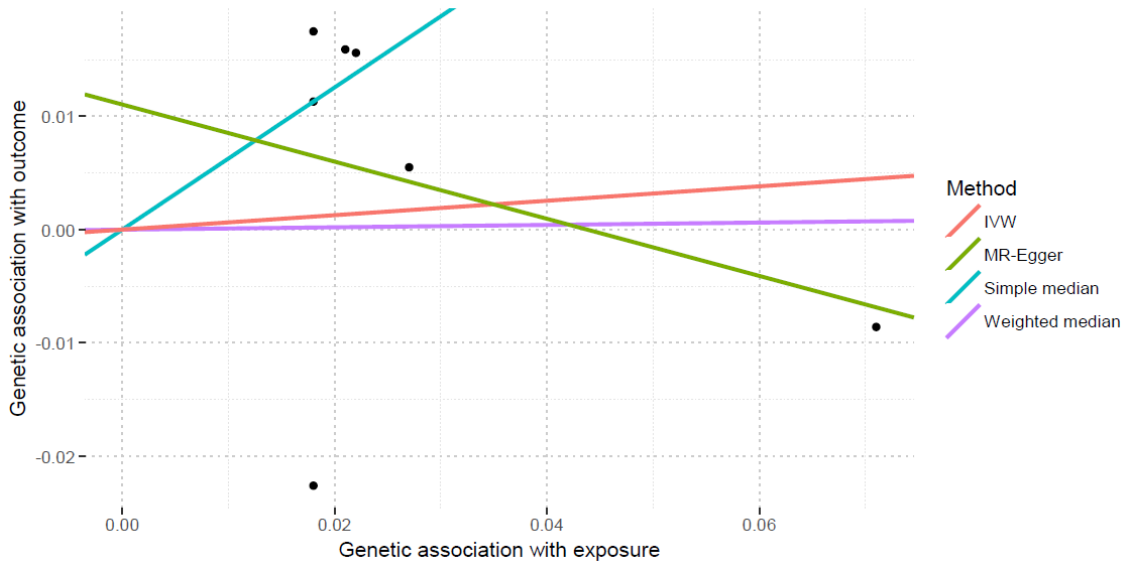

This scatter plot show individual causal estimates from each of 7 genetic variants associated with serum calcium levels on the x-axis and BMD on the y-axis. The continuous line represents the causal estimate of serum calcium levels on BMD.

**eFigure 7**. Individual genetic estimates of serum calcium levels with body-BMD in individuals 15-30 years using different methods

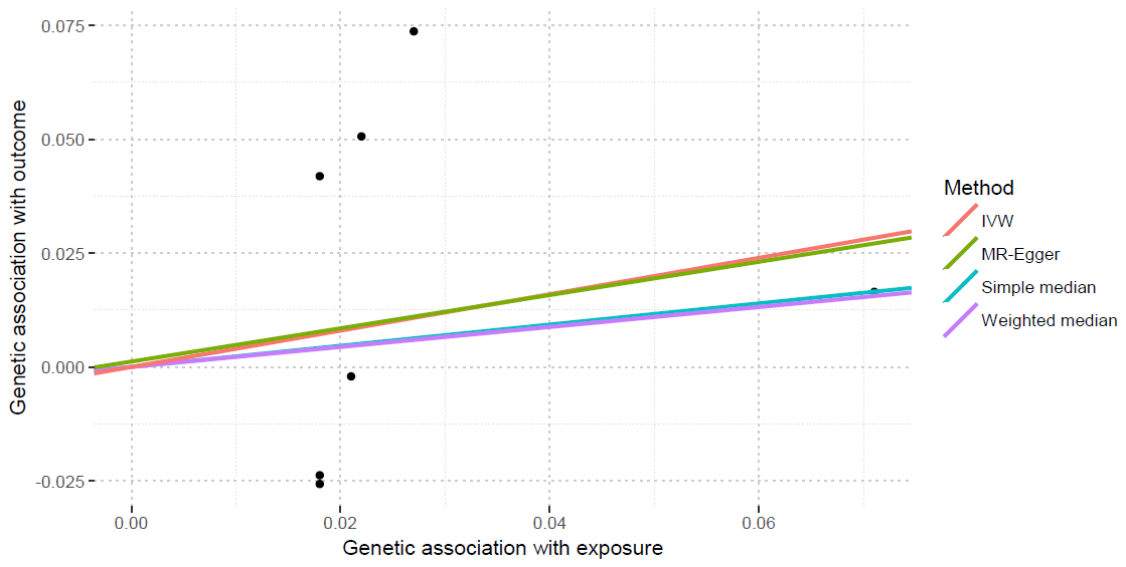

This scatter plot show individual causal estimates from each of 7 genetic variants associated with serum calcium levels on the x-axis and BMD on the y-axis. The continuous line represents the causal estimate of serum calcium levels on BMD.

**eFigure 8**. Individual genetic estimates of serum calcium levels with body-BMD in individuals 30-45 years using different methods

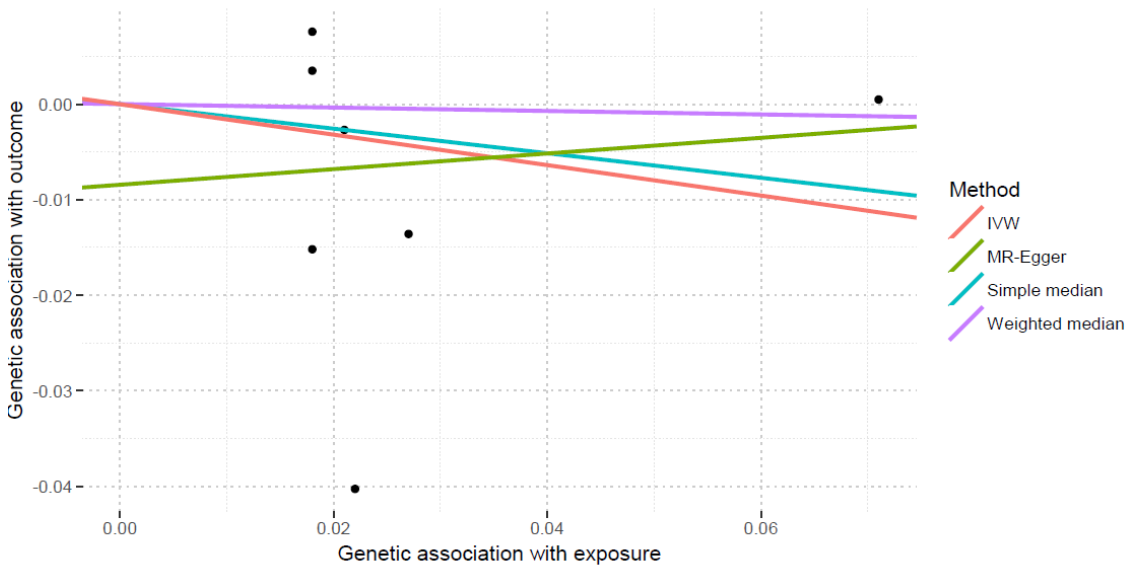

This scatter plot show individual causal estimates from each of 7 genetic variants associated with serum calcium levels on the x-axis and BMD on the y-axis. The continuous line represents the causal estimate of serum calcium levels on BMD.

**eFigure 9**. Individual genetic estimates of serum calcium levels with body-BMD in individuals 45-60 years using different methods

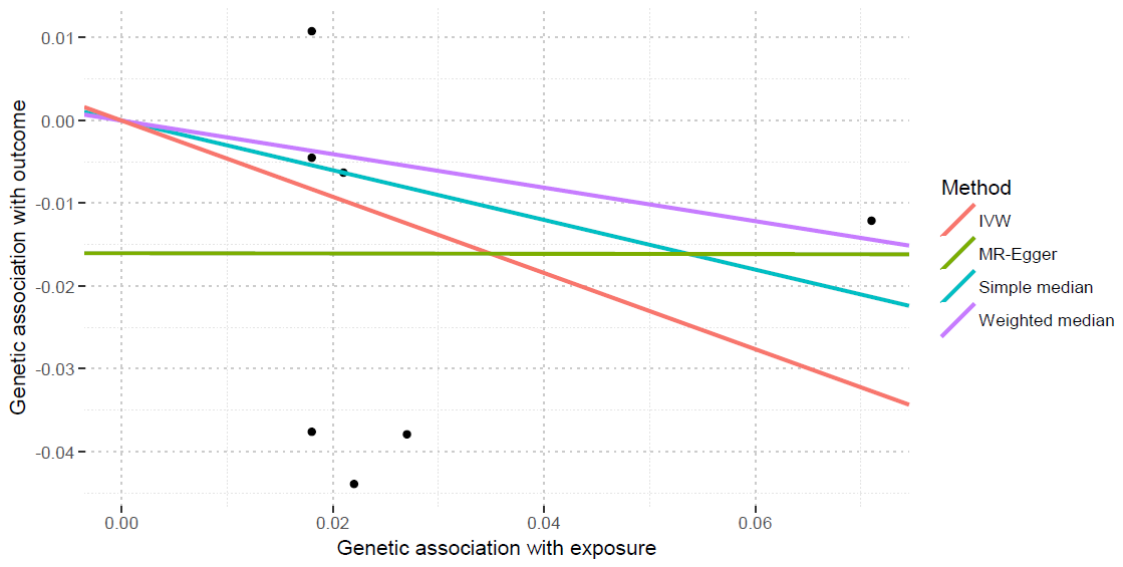

This scatter plot show individual causal estimates from each of 7 genetic variants associated with serum calcium levels on the x-axis and BMD on the y-axis. The continuous line represents the causal estimate of serum calcium levels on BMD.

**eFigure 10**. Individual genetic estimates of serum calcium levels with body-BMD in individuals 60 or more years using different methods

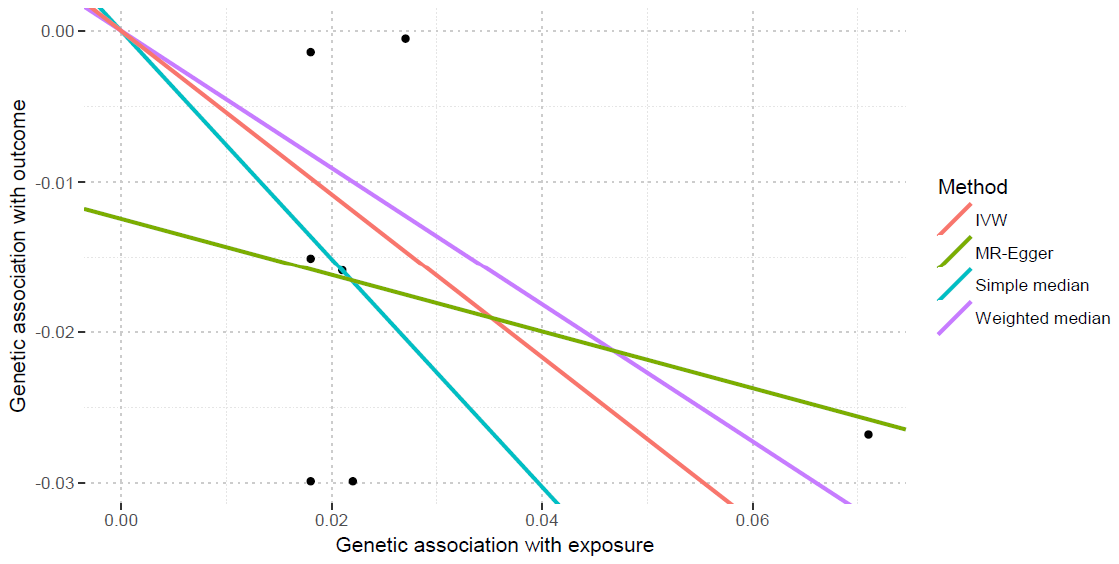

This scatter plot show individual causal estimates from each of 7 genetic variants associated with serum calcium levels on the x-axis and BMD on the y-axis. The continuous line represents the causal estimate of serum calcium levels on BMD.

**Reference**

**1.**	Cappuccio FP, Meilahn E, Zmuda JM, Cauley JA. High blood pressure and bone-mineral loss in elderly white women: a prospective study. Study of Osteoporotic Fractures Research Group. *Lancet.* Sep 18 1999;354(9183):971-975.

**2.**	Ma L, Oei L, Jiang L, et al. Association between bone mineral density and type 2 diabetes mellitus: a meta-analysis of observational studies. *Eur J Epidemiol.* May 2012;27(5):319-332.

**3.**	Shanbhogue VV, Mitchell DM, Rosen CJ, Bouxsein ML. Type 2 diabetes and the skeleton: new insights into sweet bones. *Lancet Diabetes Endocrinol.* Feb 2016;4(2):159-173.

**4.**	Lorentzon M, Mellstrom D, Haug E, Ohlsson C. Smoking is associated with lower bone mineral density and reduced cortical thickness in young men. *J Clin Endocrinol Metab.* Feb 2007;92(2):497-503.

**5.**	Law MR, Hackshaw AK. A meta-analysis of cigarette smoking, bone mineral density and risk of hip fracture: recognition of a major effect. *BMJ.* Oct 4 1997;315(7112):841-846.

**6.**	Baron JA, Farahmand BY, Weiderpass E, et al. Cigarette smoking, alcohol consumption, and risk of hip fracture in women. *Arch Intern Med.* Apr 9 2001;161(7):983-988.

**7.**	Jang HD, Hong JY, Han K, et al. Relationship between bone mineral density and alcohol intake: A nationwide health survey analysis of postmenopausal women. *PLoS One.* 2017;12(6):e0180132.

**8.**	McLernon DJ, Powell JJ, Jugdaohsingh R, Macdonald HM. Do lifestyle choices explain the effect of alcohol on bone mineral density in women around menopause? *Am J Clin Nutr.* May 2012;95(5):1261-1269.

**9.**	Lodder MC, de Jong Z, Kostense PJ, et al. Bone mineral density in patients with rheumatoid arthritis: relation between disease severity and low bone mineral density. *Ann Rheum Dis.* Dec 2004;63(12):1576-1580.

**10.**	Scott EM, Gaywood I, Scott BB. Guidelines for osteoporosis in coeliac disease and inflammatory bowel disease. British Society of Gastroenterology. *Gut.* Jan 2000;46 Suppl 1:i1-8.

**11.**	Bjarnason I, Macpherson A, Mackintosh C, Buxton-Thomas M, Forgacs I, Moniz C. Reduced bone density in patients with inflammatory bowel disease. *Gut.* Feb 1997;40(2):228-233.

**12.**	Ho SC, Chen YM, Woo JL. Educational level and osteoporosis risk in postmenopausal Chinese women. *Am J Epidemiol.* Apr 1 2005;161(7):680-690.

**13.**	Solomon DH, Avorn J, Canning CF, Wang PS. Lipid levels and bone mineral density. *Am J Med.* Dec 2005;118(12):1414.

**14.**	Mahajan A, Go MJ, Zhang W, et al. Genome-wide trans-ancestry meta-analysis provides insight into the genetic architecture of type 2 diabetes susceptibility. *Nat Genet.* Mar 2014;46(3):234-244.

**15.**	Locke AE, Kahali B, Berndt SI, et al. Genetic studies of body mass index yield new insights for obesity biology. *Nature.* Feb 12 2015;518(7538):197-206.

**16.**	Shungin D, Winkler TW, Croteau-Chonka DC, et al. New genetic loci link adipose and insulin biology to body fat distribution. *Nature.* Feb 12 2015;518(7538):187-196.

**17.**	Ehret GB, Munroe PB, Rice KM, et al. Genetic variants in novel pathways influence blood pressure and cardiovascular disease risk. *Nature.* Sep 11 2011;478(7367):103-109.

**18.**	Genome-wide meta-analyses identify multiple loci associated with smoking behavior. *Nat Genet.* May 2010;42(5):441-447.

**19.**	Schumann G, Liu C, O'Reilly P, et al. KLB is associated with alcohol drinking, and its gene product beta-Klotho is necessary for FGF21 regulation of alcohol preference. *Proc Natl Acad Sci U S A.* Dec 13 2016;113(50):14372-14377.

**20.**	Okbay A, Beauchamp JP, Fontana MA, et al. Genome-wide association study identifies 74 loci associated with educational attainment. *Nature.* May 26 2016;533(7604):539-542.

**21.**	Tillmann T, Vaucher J, Okbay A, et al. Education and coronary heart disease: mendelian randomisation study. *BMJ.* Aug 30 2017;358:j3542.

**22.**	Dale CE, Fatemifar G, Palmer TM, et al. Causal Associations of Adiposity and Body Fat Distribution With Coronary Heart Disease, Stroke Subtypes, and Type 2 Diabetes Mellitus: A Mendelian Randomization Analysis. *Circulation.* Jun 13 2017;135(24):2373-2388.

**23.**	Burgess S, Thompson SG. Interpreting findings from Mendelian randomization using the MR-Egger method. *Eur J Epidemiol.* May 2017;32(5):377-389.

**24.**	Burgess. S, Bowden. J, Dudbridge. F, Thompson. SG. Robust instrumental variable methods using multiple candidate instruments with application to Mendelian randomization. *arXiv.* May 2017.

**25.**	O'Seaghdha CM, Wu H, Yang Q, et al. Meta-analysis of genome-wide association studies identifies six new Loci for serum calcium concentrations. *PLoS Genet.* 2013;9(9):e1003796.

**26.**	Zheng HF, Forgetta V, Hsu YH, et al. Whole-genome sequencing identifies EN1 as a determinant of bone density and fracture. *Nature.* Oct 1 2015;526(7571):112-117.

**27.**	Kemp JP, Morris JA, Medina-Gomez C, et al. Identification of 153 new loci associated with heel bone mineral density and functional involvement of GPC6 in osteoporosis. *Nat Genet.* Oct 2017;49(10):1468-1475.

**28.**	Medina-Gomez C, Kemp JP, Trajanoska K, et al. Life-Course Genome-wide Association Study Meta-analysis of Total Body BMD and Assessment of Age-Specific Effects. *Am J Hum Genet.* Jan 4 2018;102(1):88-102.
